# Conserved visual capacity of rats under red light

**DOI:** 10.1101/2020.11.05.370064

**Authors:** Nader Nikbakht, Mathew E. Diamond

## Abstract

Recent studies examine the behavioral capacities of rats and mice with and without visual input, and the neuronal mechanisms underlying such capacities. These animals are assumed to be functionally blind under red light, an assumption that might originate in the fact that they are dichromats who possess ultraviolet and green but not red cones. But the inability to see red as a color does not necessarily rule out form vision based on red light absorption. We measured Long-Evans rats’ capacity for visual form discrimination under red light of various wavelength bands. Upon viewing a black and white grating, they had to distinguish between two categories of orientation, horizontal and vertical. Psychometric curves plotting judged orientation versus angle demonstrate the conserved visual capacity of rats under red light. Investigations aiming to explore rodent physiological and behavioral functions in the absence of visual input should not assume red-light blindness.

## Introduction

Rats, like many rodents, are largely crepuscular and, even during daylight, are usually to be found in poorly illuminated environments (Macdonald et al., 1994). Their retina is rod-dominated, with cones making up as little as 1% of photoreceptors (Jacobs et al., 2001; La Vail, 1976). Although rats are not color-blind (Jacobs et al., 2001; Lemmon and Anderson, 1979; Muenzinger and Reynolds, 1936; Munn and Collins, 1936; Walton and Bornemeier, 1938), they perform poorly in discriminating between nearby wavelengths, compared to humans (Walton, 1933). Color sensitivity is based on two sets of cones with peak sensitivities in the range of ultraviolet (UV cones; peak absorption at 358–359 nm) and green (M cones; peak absorption at 509–510 nm) (Deegan and Jacobs, 1993; Jacobs et al., 1991; Szél and Röhlich, 1992). Recently, electroretinogram (ERG) responses of the photopic spectral sensitivity curves of photoreceptors of rats and mice were measured throughout the UV-visible spectrum (300 to 700 nm) (Rocha et al., 2016). These measurements identified two sensitivity peaks in Wistar rats, 362 and 502 nm; no significant response to long wavelength light (above 620 nm) was detected. This study reinforced the already existing notion that red light is experienced as a total absence of usable light. In the present study we challenge the notion of form vision blindness under red light and find, contrary to expectation, good performance.

## Results

Rats were required to categorize the orientation of a solid disk-like object with a circular boundary and raised parallel bars, alternately colored white and black, thus forming a square wave grating (Figure 1A). Orientations in the range of 0°– 45° were rewarded as horizontal and orientations in the range of 45°– 90° as vertical (Figure 1B). Figure 1C illustrates the sequence of events in the behavioral task. Each trial started with the rat’s head poke, which triggered the opening of an opaque gate, followed by illumination with light sources of various wavelengths. A transparent panel in front of the object prevented the rat from generating tactile cues. After observing the object, the rat turned its head toward one spout (L or R) and licked. The boundary angle, 45°, was rewarded randomly on left or right. Illumination was by a white LED array or else by monochrome LEDs with peak intensities at 626 nm, 652 nm, 729 nm, 854 nm and 930 nm as measured by spectrometer and verified by the manufacturer’s datasheet (Figure 1D; see Methods). Half widths were 12.1–49.1 nm.

**Figure 1.**
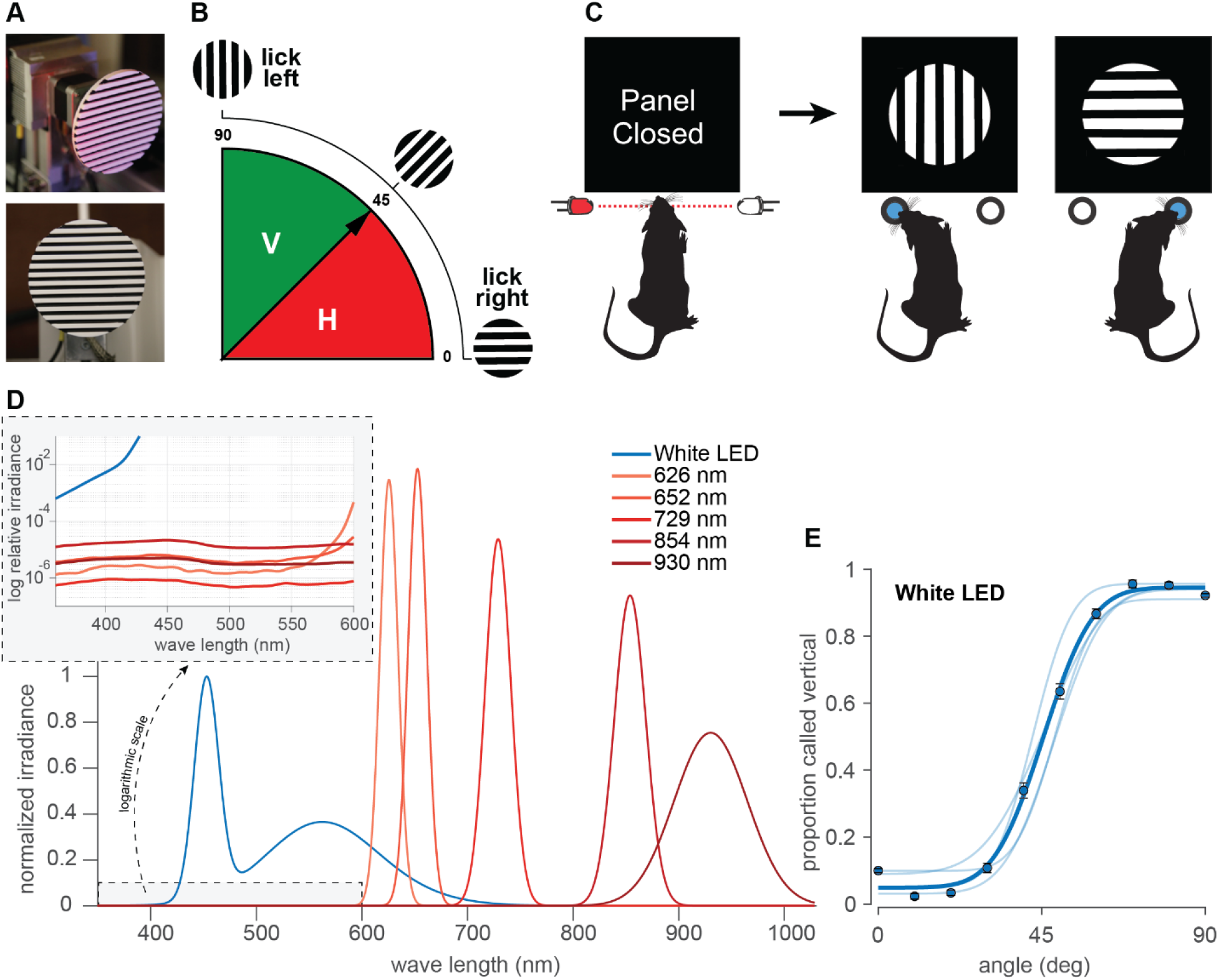
Orientation categorization task. (A) Discriminandum viewed at an oblique angle (top) and exactly from the front (bottom), the latter approximating the perspective of the rat. (B) Schematic of the orientations of the stimuli and rule of the categorization task. 0°– 45° (red) rewarded as “horizontal” and 45°– 90° (solid green) rewarded as “vertical.” (C) Sequential steps in the behavioral task. Each trial started with a head poke that interrupted a light beam and triggered the opening of an opaque gate, followed by visual access to the object. After probing the stimulus, the rat turned its head toward one spout, in this illustration left for vertical and right for horizontal. See Figure S1A for the experimental setup. (D) Gaussian fits to the normalized irradiance for each LED, measured by a spectrometer. Each distribution has been normalized to give the same area under the curve to improve visualization; non-normalized values are shown in Figure S2. The inset uses a log scale of the relative intensities to show that < 600 nm-irradiance emitted by the red LEDs was many orders of magnitude below that of the white LED. Additionally, despite ~10-30 times higher intensity of shorter wavelength emissions of the infrared LEDs (854 nm and 930 nm) compared to red LEDs, rats performed at chance under illumination with IR LEDs providing further evidence that visual performance under red light could not be well explained by unintended “leakage” towards shorter wavelengths. (E) Pale curves give the performance of 4 rats under white light. Dark data points and curves show the average over all rats. Error bars are 95% binomial confidence intervals. See Figure S1A for the experimental setup.

To quantify rats’ performance, we used a cumulative Gaussian function to fit psychometric curves to the data of each rat (see Methods). Figure 1E reveals that all rats (N = 4, pale curves; average curve in dark blue) performed well under white LED illumination.

Sessions with white light were interspersed with sessions illuminated by various narrow-band monochrome LEDs in the range of red, far-red, and infrared (Figure 1D). It is important to perform behavioral testing under illumination with narrow-band monochromatic light sources to eliminate the possibility of off-peak illumination through the tail of a wide power spectral distribution. Figure 2A-B shows that dark-adapted rats performed the visual categorization task under 626 nm and 652 nm LEDs (perceived as red by humans) with accuracy equivalent to that under white light, refuting the common belief of functional blindness under red light. Rats performed well even under peak 729 nm (perceived as far-red by humans), though clearly diminished with respect to white (Figure 2C). They performed poorly under infrared illumination, comprising 854 nm and 930 nm light (Figure 2D, E).

**Figure 2.**
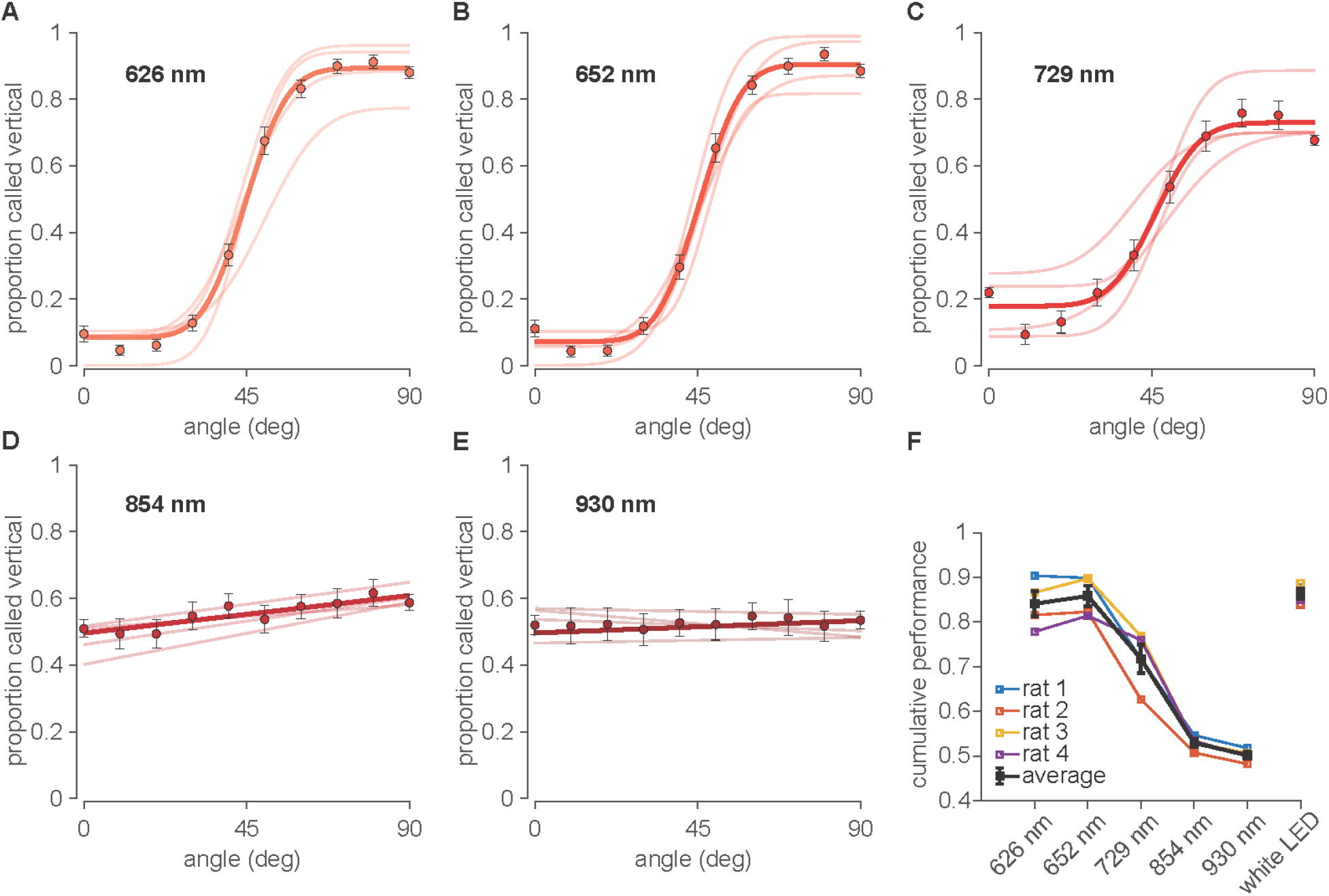
Performance under far-red and infrared illumination. (A-E) Psychometric curves obtained under illumination with monochrome LEDs in the range of far red to infrared with peak wavelengths at 626 nm, 652 nm, 729nm, 854nm and 930nm, respectively. Pale curves depict the performance of 4 rats. Dark data points and lines or curves show the average over all rats. Error bars are 95% binomial confidence intervals. (F) Summary cumulative performance (proportion correct) for all 4 rats (colored) and the average rat (black) in each illumination condition. Data from all angles (except 45°, uninformative about discriminative capacity) are pooled. See Figure S1B for tests of significance.

The performance (N=4 rats) under different illumination wavelengths is summarized by the cumulative proportion of trials categorized correctly across all orientations (Figure 2F). Average performance was 87±2% correct under white light (confidence interval is SEM across rats). Average performance was 84±3% under 626 nm illumination (p=0.158 compared to white light; bootstrap tests illustrated in Supplementary Figure S1B), 86±2% under 652 nm (p=0.419 compared to white light), 72±3% under 729 nm (p=0.00 compared to white light), 53±2% under 854 nm (p=0.257 compared to chance) and 50±1% under 930 nm average performance (p=0.316 compared to chance).

## Discussion

Red illumination is often used in reverse light cycle conditions in animal husbandry settings with the assumption that it will not affect rodent circadian rhythms (Emmer et al., 2018). Furthermore, experiments done under red light are believed to place the animal in dark conditions while allowing the experimenter to observe the preparation directly or by video recording (Cloke et al., 2015; Englund et al., 2020; van Goethem et al., 2012; Harris and Diamond, 2000; Harris et al., 1999; Jacklin et al., 2016; Nikbakht et al., 2012; Pacchiarini et al., 2020; Reid et al., 2014; Salaberry et al., 2017; Sieben et al., 2015; Vasconcelos et al., 2011; Winters and Reid, 2010). But from the inability to see red as a color, it does not necessarily follow that rodents cannot absorb red light through their rod-dominated retina to support form vision. A recent study (Niklaus et al., 2020) examined the retinal responses mediated by rods and cones of pigmented (Brown Norway) and albino (Wistar) rats in response to monochromatic far red light of 656 ± 10 nm in both photopic (light-adapted) and dark-adapted (scotopic) settings. Both rat strains showed significant scotopic and photopic ERG responses to red light even at low intensities. These results hinted that even far-red light may provide effective illumination for rats. However, whether the photoreceptor activation by red light can lead to functionally meaningful signals has not yet been established. Excitation of retinal receptors by red light does not, by itself, indicate behavioral availability of the signal. For instance, red light might act to entrain circadian rhythms, unconscious to the animal.

Though the photoreceptor mechanisms that underlie the surprising form vision under red light are beyond the scope of this report, it is interesting to note the possibility that cone photoreceptors could respond to longer wavelengths through a nonlinear optical process including two-photon activation of rhodopsins (Palczewska et al., 2014; Vinberg et al., 2019).

In the present work, Long-Evans rats demonstrated substantial visual form capacity under red and far-red light. Object discrimination began to degrade when illumination wavelength increased from 652 to 729 nm and was almost nil at 854 nm. The performance curves suggest the optimal conditions for vision-excluded behavioral studies. Illumination at 850 nm did not support visual form capacity, yet remains within the sensitivity range of inexpensive silicone detectors (CMOS and CCDs). Thus, behaviors may be documented by video while the animal performs without visual cues.

Rats and mice are the most frequently used laboratory mammals, ideal for research on spatial navigation (Frank et al., 2000; MacDonald et al., 2011; Moser et al., 2017; O'keefe and Nadel, 1978; Pastalkova et al., 2008; Wood et al., 2000) and the processing of tactile (Diamond et al., 2008; Fassihi et al., 2017; Zuo and Diamond, 2019) and olfactory (Chae et al., 2019; Koldaeva et al., 2019; Uchida and Mainen, 2003; Uchida et al., 2006) information. Neuroscientists have not traditionally attributed to rodents the wide range of visual perceptual functions characteristic of primates. However there is growing interest in the use of rodents for the study of vision, alone or combined with other modalities (Gharaei et al., 2018; Nikbakht et al., 2018; Nikbakht Nasrabadi, 2015; Sieben et al., 2015; Zoccolan, 2015). Rats under broad-wavelength conditions spontaneously recognize an object even when views differ by angle, size, and position (Zoccolan, 2015); such generalization is a hallmark of authentic visual perception and was once believed to belong only to primates. Importantly, they achieve high level sensory-perceptual cognition through the workings of neuronal circuits that are accessible (Crochet et al., 2019; Matteucci and Zoccolan, 2020; Matteucci et al., 2019; Steinmetz et al., 2019; Tafazoli et al., 2017). The present study extends the range of visual perceptual functions for which rats can serve as models, characterizing their performance in judging bar orientation and the longest illumination wavelengths at which this capacity remains intact. A complete understanding of the visual processing of these animals is important not only in the design and control of the behavioral and physiological experiments but also to ensure optimal environmental lighting conditions for their well-being in laboratory settings.

## Materials and methods

### Experimental Subject Details

Four male Long–Evans rats (Charles River Laboratories, Calco, Italy) were used. They were caged in pairs and maintained on a 12/12 hr light/dark cycle; experiments were conducted during the light phase. Upon arrival they were 8 weeks old, weighing approximately 250 g, and typically grew to over 600 g over the course of the study. They had free access to food in the cage. To promote motivation in the behavioral task, rats were water-deprived on days of training/testing. During each session they received 17-22 mL of pear juice diluted in water (1 unit juice: 4 units water) as reward. After the session they were given access to water ad libitum for 1 hr, though they were rarely thirsty then they were placed for several hours in a large, multistory enriched environment to interact with other rats. Animals were examined weekly by a veterinarian. Protocols conformed to international norms and were approved by the Ethics Committee of SISSA and by the Italian Health Ministry (license numbers 569/2015-PR and 570/2015-PR).

### Behavioral Method Details

#### Apparatus

The main chamber of the apparatus, custom-built in opaque white Plexiglas, measured 25 × 25 × 37 (H × W × L, cm) (Figure S1A). The rat started a trial by interrupting an infrared beam detected by a phototransistor (Figure 1C). Beam interruption triggered fast opening of an opaque panel (through a rotational motion of 40° in 75 ms), actuated by a stepper-motor, uncovering a circular hole (diameter 5 cm) in the front wall through which the rat could extend its head to see the object. The stimulus was 3 cm behind the opaque panel (further details below) and the reward spouts were 2 cm lateral to the edge of the stimulus. A transparent panel prevented direct touch.

The apparatus was in a Faraday cage which, with the door closed, provided acoustic, visual, and electromagnetic isolation. An array of 12 infrared emitters (λ = 930 nm, OSRAM Opto Semiconductors GmbH, Germany) illuminated the stimulus port to permit the investigator to monitor behavior and to execute video recording. Such illumination did not provide visual cues for the rat (see Results). For visual testing different light sources were used to illuminate the stimulus: a pair of 6 white LED arrays or various high power monochrome LED arrays (Roithner LaserTechnik, GmbH) with peak intensities at 626 nm (LED620-66-60), 652nm (LED660-66-60), 729 nm (LED735-66-60), 854 nm (LED850-66-60) and 930 nm (LED940-66-60) (see Figure 1D for gaussian fits to the measured spectrum in ambient temperature). The power spectral distributions were measured with a calibrated spectrometer and the associated software (Model: FLAME-S-XR1-ES, OceanOptics, Rochester, NY) via a visible-NIR fiber (core diameter 200 μm) attached to a cosine corrector. The cosine corrector acts as an optical diffuser that couples to the optical fiber and spectrometer to collect signals from 180° field of view.

Two infrared-sensitive video systems (Point Grey Flea, Edmund Optics, Barrington, NJ) registered the rat’s actions. The first camera, equipped with a macro lens (Fujinon TV HF25HA-1B Lens, Fujifilm, Tokyo) mounted 25 cm above the stimulus delivery area (distance with respect to the center of stimulus), monitored the rat’s interaction with the object. In some sessions, this camera was set to 250 f/s to monitor head, snout and whisker position and movement during behavior. The second camera provided a wide-angle view (Fujinon HF9HA-1B Lens, Fujifilm, Tokyo) and monitored the entire setup, illuminated with adjustable infrared LEDs, at 30 f/s.

Reward spouts included custom-made infrared diode sensors interrupted by the tongue. Only the licking signal from the correct spout triggered the pump motor (NE-500 programmable OEM; New Era Pump Systems, mounted on a vibration-cancellation pedestal) to extrude the reward, 0.05 mL per trial of diluted pear juice. Licking marked the end of the trial, accompanied by the closure of the opaque front panel. Before the next trial began, the motor on which the stimulus was mounted rotated to generate the next orientation.

Custom made software was developed using LabVIEW (National Instruments, Austin, TX). An AVR32 board (National Instruments) and multiple Arduino Shields (National Instruments) acquired all sensor signals and controlled the motors, LEDs, and the reward syringe pumps. All the sensors, actuators (including motors and pumps) and lights were interfaced with the computer program allowing full control over a wide range of parameters governing the flow of the training and testing. Although fully automatic, the software allowed the experimenter to modify all the parameters of the task and control the lights, sensors and motors online as needed.

#### Visual Stimulus Presentation

The stimulus was a black and white square-wave grating within a circular 9.8 cm-diameter circumference, built in-house by a 3D printer (3D Touch, BFB Technologies, Figure 1A). It was mounted on a stepper-motor and rotated to generate the trial’s intended orientation (Figure 1B).

The stimulus stepper-motor was controlled through a feedback system with a digital step counter to maintain the exact desired orientation. In the study that originally explored orientation judgment (Nikbakht et al., 2018), tactile exploration of the object was allowed in some trials, but the tactile condition is not considered in the present work; only visual, touch-free data are included. Within behavioral testing sessions each stimulus orientation was sampled from a uniform distribution in 5° steps between −45° and 135° and presented in a semi-random fashion (sampling without replacement). For analysis we binned the angles every 10°. Visual acuity is measured in cycles per degree (cpd), an assessment of the number of lines that can be seen as distinct within a degree of the visual field:

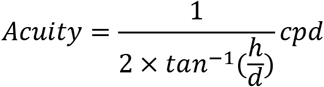

where *h* is the width of each line in the stimulus and *d* is the distance from the eye. Considering the 3 cm distance behind the opaque panel (Figure S1A), at the moment of panel opening each cycle of the 14 cycles would occupy 2 × *tan* – 1(3.5/30) = 13.31° of visual angle, for a total stimulus coverage of about 117°. The spatial frequency of the gratings would be 1/(2 × *tan* – 1(3.5/30)) = 0.075 cycles per degree of visual angle. As the normal visual acuity of Long–Evans rats has been estimated as ~1 cpd (Prusky et al., 2002), the bars would be expected to be resolvable. Rats also have a large depth of focus, from 7 cm to infinity (Powers and Green, 1978). The width of the binocular field directly in front of the rat’s nose, generally considered the animal’s binocular viewing area (Mei et al., 2012), ranges from approximately 40°-110°, depending on head pitch (Wallace et al., 2013). The 117° stimulus should thus completely cover the rat’s binocular visual field.

When all illuminations were off, the ambient light magnitude was 0 cd/mm2 (Konica Minolta LS-100 luminance meter, Tokyo).

#### Behavioral Task and Training

Duration of training to reach stable performance was typically 4–6 weeks, with 1 session per day, and varied according to individual differences in rate of learning. The standard training was done with stimulus illumination by white LEDs. The training protocol proceeded across a sequence of stages given in Nikbakht et al. (2018). Exclusion of unintended stimulus cues (olfaction, rotating motor noise) was controlled as given in (Nikbakht et al., 2018). Rats were dark-adapted in a light-free environment for 20-30 minutes prior to each session.

### Quantification and Statistical Analysis

#### Analysis of Behavioral Data

We analyzed the behavioral data in MATLAB (MathWorks, Natick, MA) and LabVIEW. To quantify a single rat’s performance, we fit psychometric curves to its choice data. For a given orientation, we calculated the proportion of trials categorized as vertical. Ideally, rats would categorize all trials with angle greater than 45° as vertical and all trials with angle less than 45° as horizontal. For 45° trials, choices should be evenly distributed between vertical and horizontal. However, task difficulty grows in the vicinity of 45°, such that real performance is better described by a sigmoid function with an inflection point at the point of subjective equality (PSE), the orientation at which subjects report the stimulus with equal likelihood as horizontal or vertical. In unbiased rats, the PSE should be at 45°. We generated psychometric functions using a cumulative Gaussian function with the general form given in the equation below based on (Wichmann and Hill, 2001). The parameter estimation was then performed in MATLAB using maximum-likelihood estimation:

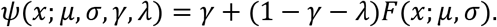

The two-parameter function *F*(*x*;*μ,σ*), is defined by a cumulative Gaussian distribution, as follows:

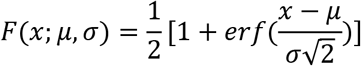

where *x* is the stimulus orientation, *γ* is the lower-bound of the function ‡, and *λ* is the lapse rate. Often, *γ* and *λ* are considered to arise from stimulus-independent mechanisms of guessing and lapsing. *μ* is the mean of the probability distribution that determines the displacement along the abscissa of the psychometric function–a reflection of the subject’s bias–and *σ* is the standard deviation of the cumulative Gaussian distribution. *σ* determines the slope of the psychometric function, a common measure of acuity.

Where fitting with a sigmoid was not appropriate (Figure 2 D-E), data were fit with the line *y* = *ax* + *y*_0_, where *x* is the stimulus orientation, *a* is the slope and *y*_0_ is the y-intercept.

#### Test of Significance for the Fitted Psychometric Curves

Errors around the performance value for each orientation and modality condition were expressed as a 95% binomial proportion confidence interval computed by approximating the distribution of errors about a binomially-distributed observation, 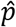, with a normal distribution:

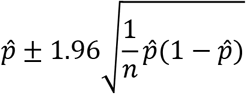

where 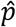 is the proportion of correct trials (Bernoulli) and *n* is the number of trials.

For statistical tests of significance, we performed a non-parametric test based on bootstrapping, as follows. We computed a distribution of the performance values from the fitted psychometric functions based on 1,000 resamples of the behavioral data. We then performed pairwise comparisons between all the performance values generated via bootstrapping from fitted psychometric functions of each experimental condition, calculated the overlap between the distributions and computed the p-values.

## Acknowledgments

We acknowledge the financial support of the Human Frontier Science Program (M.E.D.) (http://www.hfsp.org; project RGP0015/2013), the European Research Council advanced grant CONCEPT (M.E.D.) (http://erc.europa.eu; project 294498). The Regional laboratory for advanced mechatronics, LAMA FVG (http://lamafvg.it) supported the design and construction of custom instrumentation. The funders had no role in study design, data collection and analysis, decision to publish, or preparation of the manuscript. Erik Zorzin assisted in spectral measurements. We are grateful to members of the Diamond lab for fruitful comments and discussions.

## Author contributions

Conceptualization, N.N., M.E.D.; Methodology, N.N., M.E.D.; Investigation – Animal Subject Training and Testing, N.N.; Formal Analysis, N.N., M.E.D.; Resources, M.E.D.; Writing N.N. and M.E.D.; Visualization, N.N.; Supervision, M.E.D.; Project Administration, M.E.D; Funding Acquisition, M.E.D.

## Supplemental Figures

**Figure S1.**
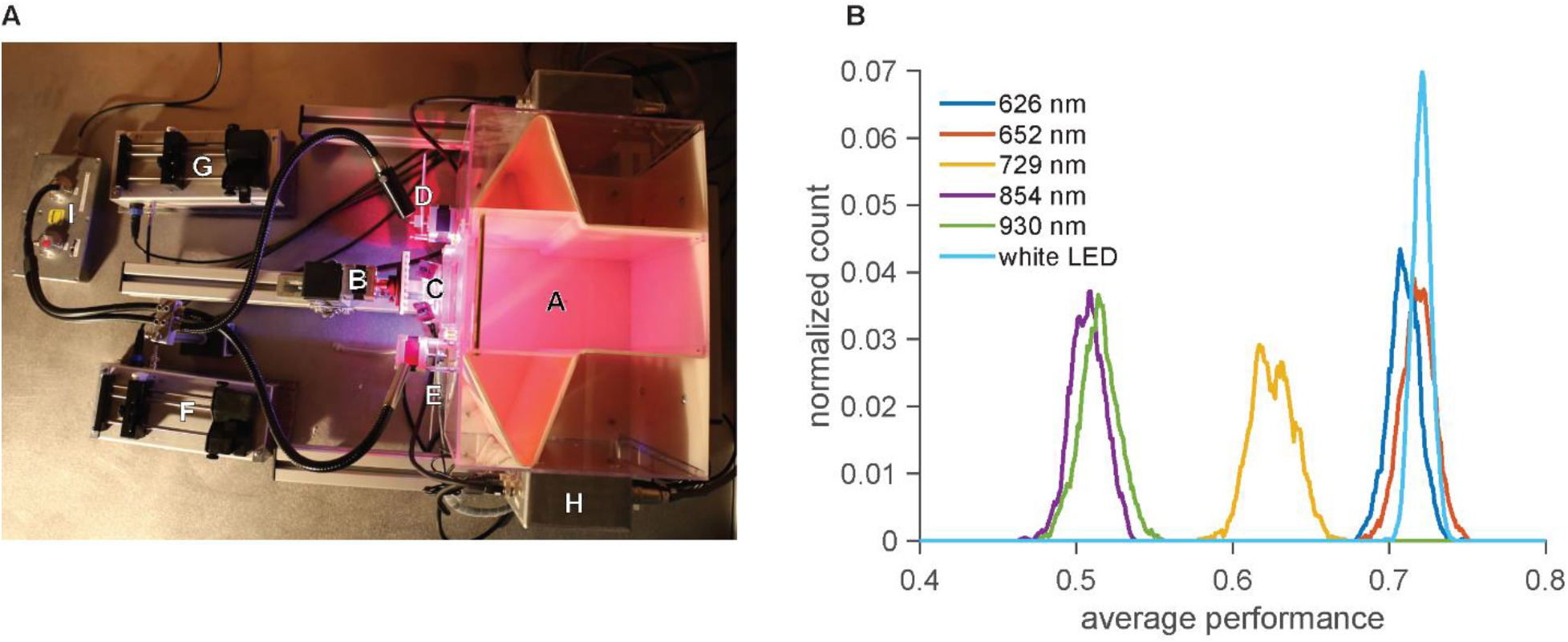
Details of the experimental setup and statistical test of significance, related to Figures 1 and 2. (A) View of the behavioral apparatus from above. Labels on the photo refer to the following components. A: the main chamber, B: stimulus and stepper motor controller together with digital step counter, C: the reward delivery area and licking sensors, D and E: transparent and opaque panels respectively, F and G: the pumps which drive the syringes loaded with diluted juice, H: the electronic control box, housing the microcontroller-based D/A boards and sensors and lights controllers, I: the light source attached to optic fibers. (B) Normalized bootstrap distributions for average illumination condition for all rats averaged performance based on 1,000 rounds of resampling in each together.

**Figure S2.**
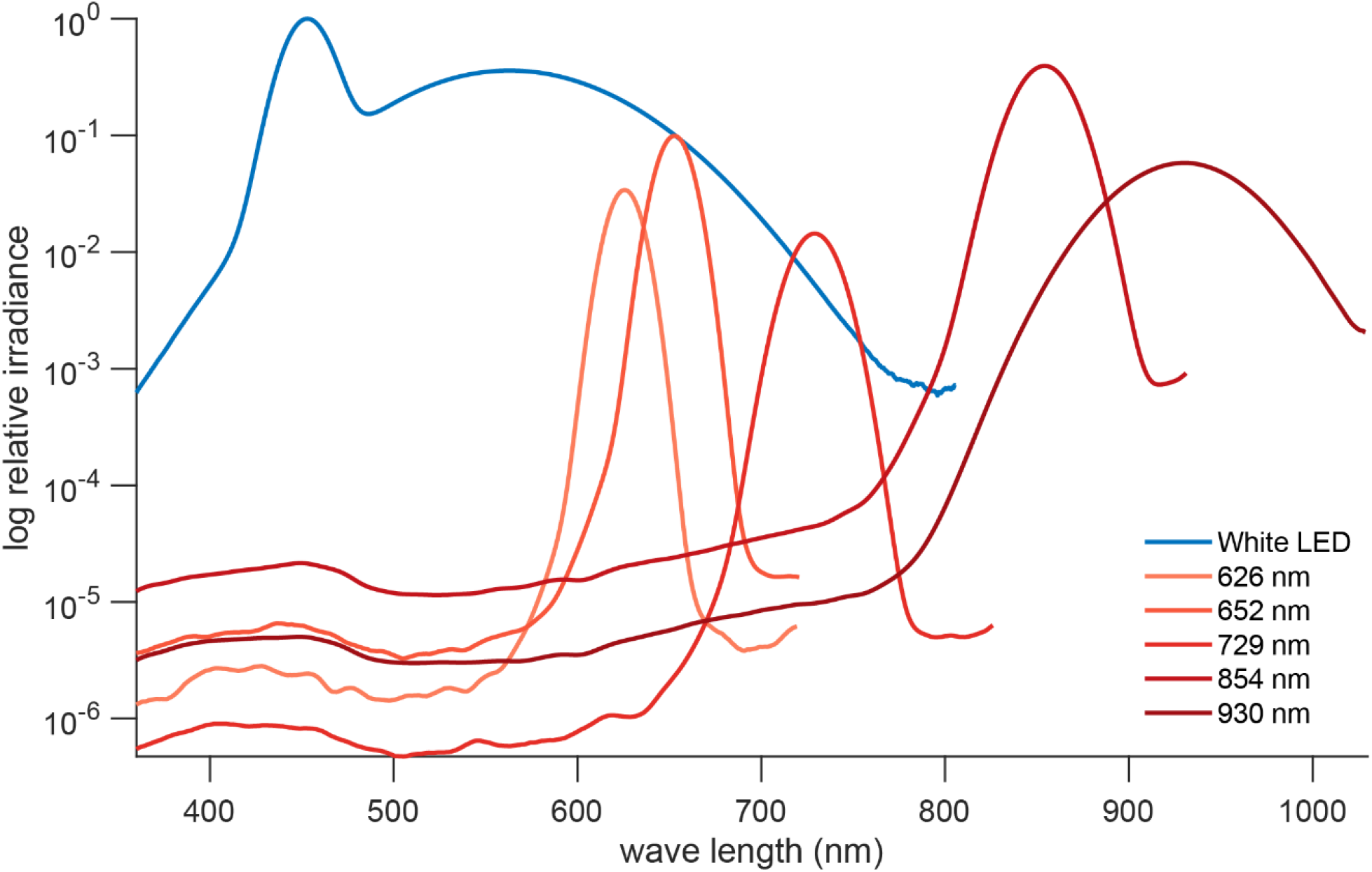
Log-scaled irradiance for each LED, measured by a spectrometer, related to Figures 1 and 2. Distributions are plotted relative to the intensity of the white LED (blue curve).

